## Supplementary Figures for "Atomistic Modeling of Liquid-Liquid Phase Equilibrium Explains Dependence of Critical Temperature on γ-Crystallin Sequence"

*Supplementary Information*

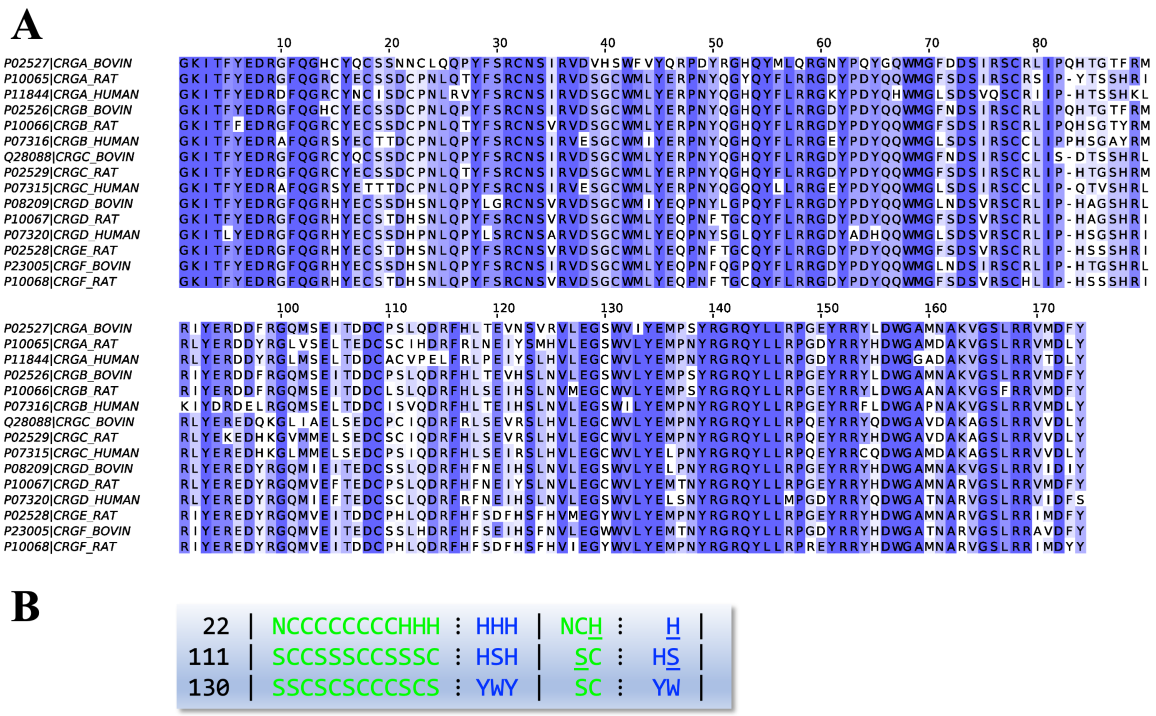


**Fig. S1.** Sequence alignment of γ-crystallins. (A) The alignment of all known sequences of bovine, human, and rat γ-crystallins by ClustalW and visualized using Jalview (https://www.jalview.org/). Residue numbers are according to bovine γB. (B) Sequence comparison at three positions showing the largest Grantham distances between the low and high-$T_c$ γ-crystallins (Fig. 5). Columns 2 and 3 display the amino acids at the same position for the 12 low-$T_c$ and 3 high-$T_c$ γ-crystallins. Columns 4 and 5 display a nonredundant list of amino acids within the low-$T_c$ and high-$T_c$ groups; amino acids that occur in both groups are underlined.

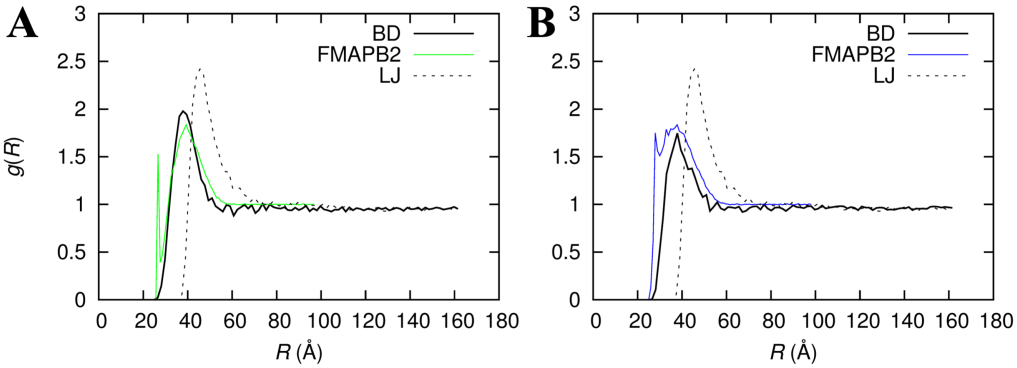


**Fig. S2.** Comparison of pair distribution functions. (A) γB. (B) γF. The "BD" trace is by counting pairs in distance bins in BD simulations at 300 K and the lowest concentration (31 mg/ml). The FMAPB2 trace is from averaging the Boltzmann factor of the pair interaction energy at 298 K in distance bins and equating the average Boltzmann factor to $g(R)$. The "LJ" trace is from counting pairs in distance bins in simulations of Lennard-Jones particles at $k_B T/\varepsilon = 1$ and $N = 30$ reported in our 2016 study. The Lennard-Jones $\sigma$ parameter was scaled to 40.5 Å, which is close to the diameter of a sphere with the steric volume $V_{st} \equiv B_2^{st}/4$ of γB.

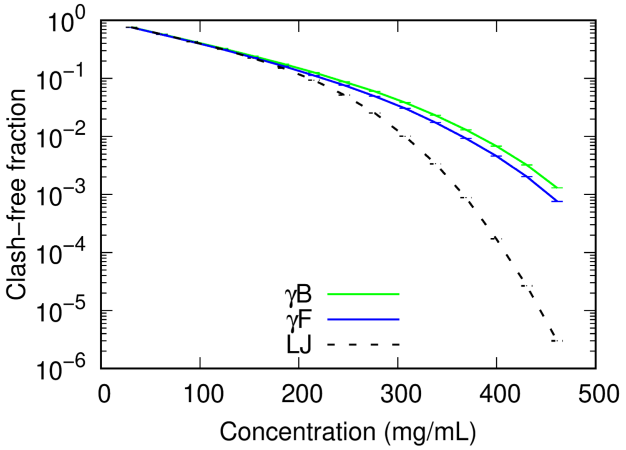


**Fig. S3.** The clash-free fraction when inserting a test protein into a protein solution. The γB and γF results were calculated by FMAPμ and averaged over 2,000 BD configurations and 500 test-protein orientations. The trace labeled "LJ" was calculated similarly on configurations generated by replacing Lennard-Jones particles with an all-atom structure of γB at an arbitrary orientation, as reported in our 2016 study (10 test-protein orientations). The simulations of Lennard-Jones particles were done at a reduced temperature $k_B T/\varepsilon = 1$ and at the same total volume and particle numbers ($N = 30$ to 450) as the BD simulations of γB and γF. Error bars were estimated by the blocking method.

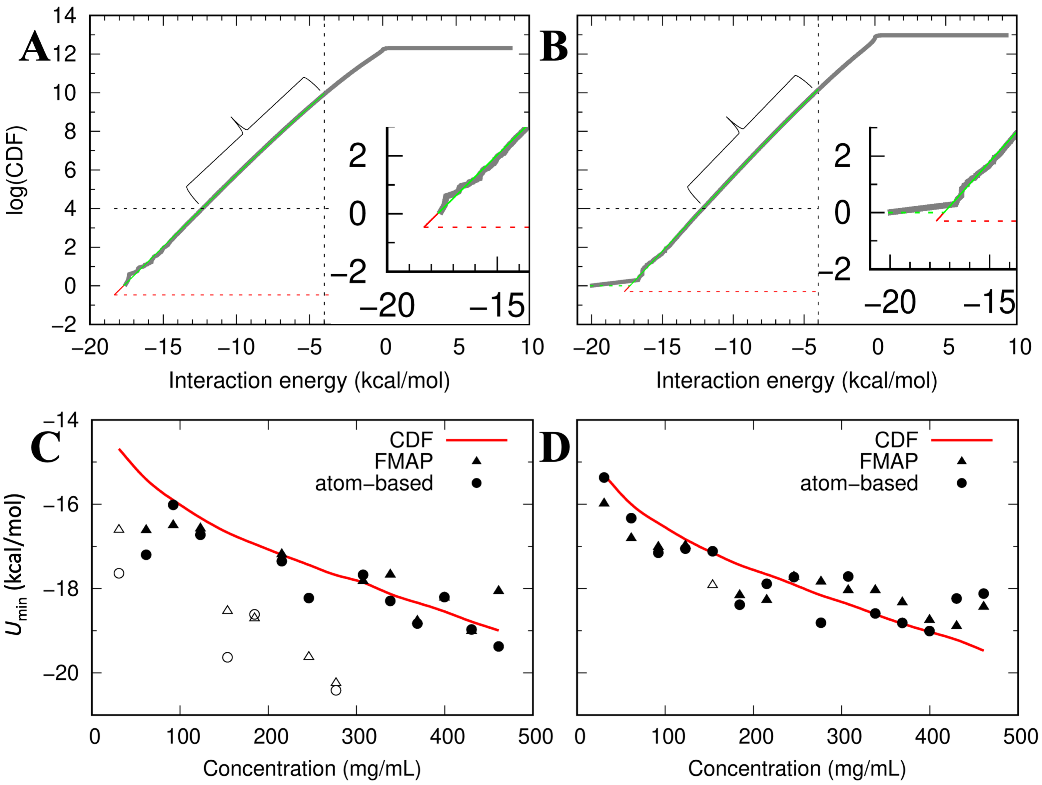


**Fig. S4.** Correction of the chemical potential by modeling the cumulative distribution function (CDF) in the low interaction energy region. (A) CDF collected from the allowed insertions in FMAPμ calculations over 2,000 BD configurations and 500 test-protein orientations, shown as gray curve for γB at 369 mg/ml. The bracketed portion, bounded from below at CDF = $10^4$ (black horizontal line) and from above at $U$ = -4 kcal/mol (black vertical line), was locally fit to a linear function. The fit function is extrapolated to the lower bound given by Eq (18) (red horizontal line). The extrapolation is shown in green down to CDF = 1 and in red beyond. A zoomed version of the lower left corner is shown in the inset. (B) Corresponding results for γB at 277 mg/ml. In contrast to (A) where the raw CDF stays close to the extrapolation all the way down to CDF = 1, a single configuration with an outlying low interaction energy (= -20.1 kcal/mol) markedly shifts the raw CDF away from the extrapolation. (C) Minimum interaction energies of γB at various concentrations. Symbols display results obtained according to Eq (17), illustrated in Fig. S5; triangles are from FMAPμ calculations whereas circles from atom-based calculations. Open symbols are for cases with an outlying lowest interaction energy, identified by a gap of at least 1.0 kcal/mol between that energy and the one from extrapolating the CDF fit function to CDF = 1, as illustrated in the panel (B) inset. Filled symbols are for cases without such outliers. The curve displays the results predicted by Eq (18). (D) The corresponding results for $\gamma F$.

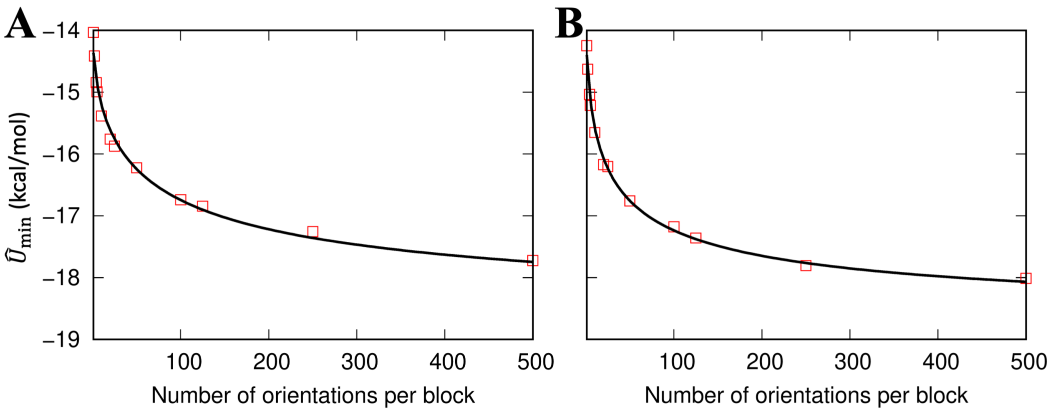


**Fig. S5.** The dependence of the mean lowest interaction energy on the block size, i.e., the number of test-protein orientations per block. Results are for (A) γB and (B) γF at 369 mg/ml. The symbols display results from the atom-based method; the curves display the fit to Eq (17).

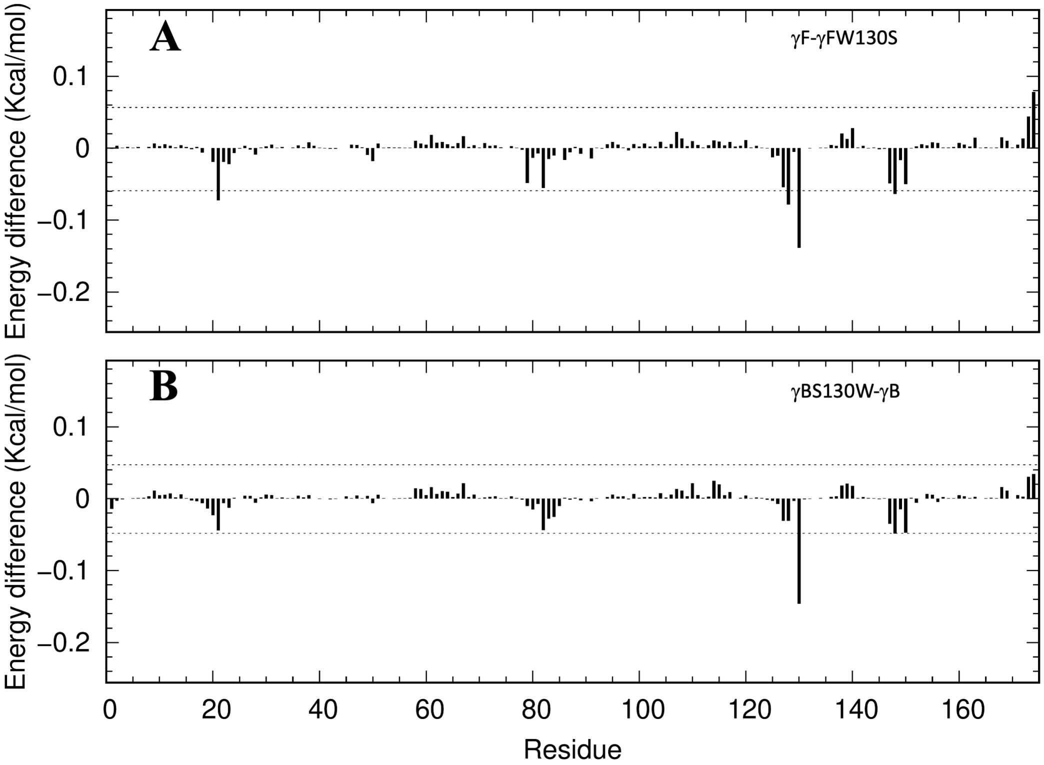


**Fig. S6.** Difference in residue-specific decomposed pair interaction energies. (A) The difference with the γFW130S results subtracted from the γF results. (B) The difference with the γB results subtracted from the γBS130W results.

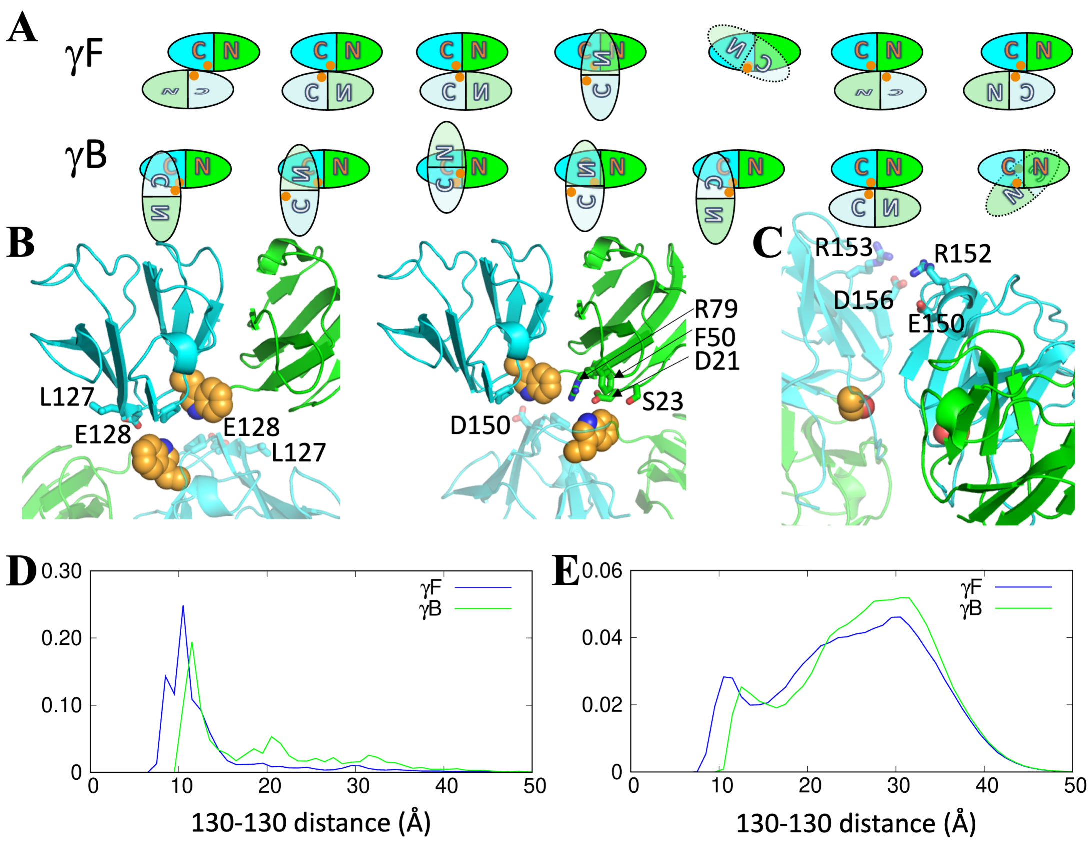


**Fig. S7.** Different tendencies of γF Trp130 and γB Ser130 to be buried in binary interfaces. (A) Illustration of the arrangements between monomers in the seven large-cluster representations of γF or γB. "C" and "N" label the N and C-terminal domains; reflected, slanted, and upside-down letters indicate self-rotation of a monomer. Orange circles represent Trp130 or Ser130 residues. (B) Cross-interface contacts in the representative poses of large cluster 1 and large cluster 3 of γF. (C) Cross-interface contacts in the representative pose of large cluster 1 of γB. (D) Distribution functions of Trp130-Trp130 distances in γF pairs (blue curve) and Ser130-Ser130 distances in γB pairs (green curve). Inter-residue distances were calculated for all poses with interaction energies < -6 kcal/mol, each weighted by the Mayer function ($T$ = 25 °C). (E) Distributions of 130-130 distances in BD simulations of γF and γB. In each snapshot, one 130-130 distance was obtained for each protein molecule, i.e., the shortest one from a neighboring molecule. The distribution function of 130-130 distances was then averaged over 2000 snapshots from γF or γB simulations at $N = 390$ (corresponding to 400 mg/ml).

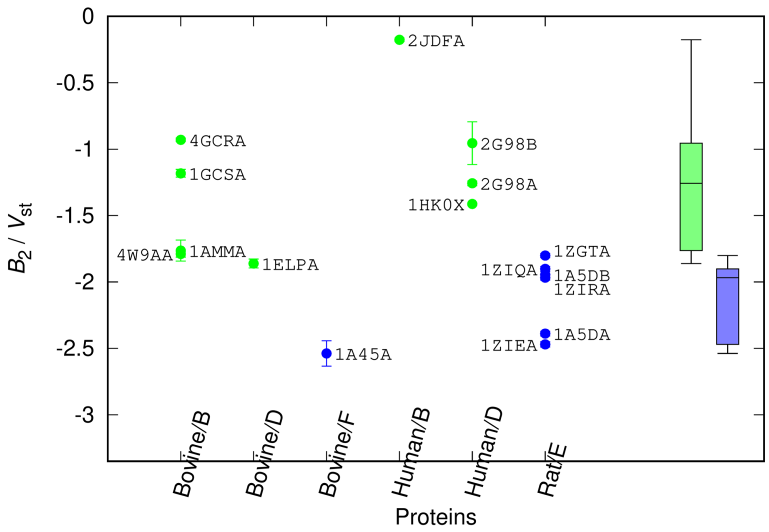


**Fig. S8.** The second virial coefficients calculated by FMAPB2 on 16 γ-crystallin structures in the PDB. The labels are the PDB entry names, with the fifth letter denoting the chain label; low and high-$T_c$ entries are in green and blue, respectively. Error bars are determined by the blocking method. Box plots for the 9 low-$T_c$ $B_2$s and 6 high-$T_c$ $B_2$s are shown at the far right.
